# Binding of a Newly Identified Essential Light Chain to Expressed *Plasmodium falciparum* Class XIV Myosin Enhances Actin Motility

**DOI:** 10.1101/127118

**Authors:** Carol S. Bookwalter, Chwen L. Tay, Rama McCrorie, Michael J. Previs, Elena B. Krementsova, Patricia M. Fagnant, Jake Baum, Kathleen M. Trybus

## Abstract

Motility of the apicomplexan parasite *Plasmodium falciparum*, the causative agent of malaria, is enabled by the glideosome, a multi-protein complex containing the class XIV myosin motor, PfMyoA. Parasite motility is necessary for invasion into host cells and for virulence. Here we show that milligram quantities of functional PfMyoA can be expressed using the baculovirus/*Sf*9 cell expression system, provided that a UCS (UNC-45/CRO1/She4p) family myosin co-chaperone from *Plasmodium spp*. is co-expressed with the heavy chain. The homologous chaperone from the apicomplexan *Toxoplasma gondii* does not functionally substitute. We expressed a functional full-length PfMyoA with bound <u>m</u>yosin <u>t</u>ail <u>i</u>nteracting <u>p</u>rotein (MTIP), the only known light chain of PfMyoA. We then identified an additional “essential” light chain (PfELC) that co-purified with PfMyoA isolated from parasite lysates. PfMyoA expressed with both light chains moved actin at ~3.8 μm/sec, more than twice that of PfMyoA-MTIP (~1.7 μm/sec), consistent with the light chain binding domain acting as a lever arm to amplify nucleotide-dependent motions in the motor domain. Surprisingly, PfMyoA moved skeletal actin or expressed *P. falciparum* actin at the same speed. Duty ratio estimates suggest that PfMyoA may be able to move actin at maximal speed with as few as 6 motors. Under unloaded conditions, neither phosphorylation of Ser19 of the heavy chain, phosphorylation of several Ser residues in the N-terminal extension of MTIP, or calcium affected the speed of actin motion. These studies provide the essential framework for targeting the glideosome as a potential drug target to inhibit invasion by the malaria parasite.

**Significance:** Motility of the apicomplexan parasite *Plasmodium falciparum*, the causative agent of malaria, relies on a divergent actomyosin system powered by the class XIV myosin, PfMyoA. We show that functional PfMyoA can be expressed in *Sf9* cells if a *Plasmodium spp*. myosin chaperone is co-expressed. We identified an “essential” light chain (PfELC) that binds to PfMyoA in parasites. *In vitro* expression of PfMyoA heavy chain with PfELC and the known light chain MTIP produced the fastest speeds of actin movement (~3.8 μm/sec). Duty ratio estimates suggest that ~6 PfMyoA motors can move actin at maximal speed, a feature that may facilitate interaction with short, dynamic *Plasmodium* actin filaments. Our findings enable drug screening for myosin-based inhibitors of *Plasmodium* cellular invasion.

## Introduction

Malaria is a blood-borne disease that causes nearly half of a million deaths per year (1), and is caused by apicomplexan parasites of the genus *Plasmodium*. The force required for the parasite to invade vertebrate host hepatocytes or erythrocytes is powered by a multi-protein assembly called the glideosome, the core of which is the class XIV myosin PfMyoA, which interacts with a divergent and dynamic *Plasmodium* actin isoform (PfACT1) (reviewed in (2)). Class XIV myosins are monomeric with a very short heavy chain. The motor domain containing the actin-binding and active site is followed by a neck region of only ~50 amino acids, and no further tail. Typical myosins have well-defined IQ motifs (consensus sequence IQxxxRGxxxR) following the motor domain, which are known to bind EF-hand proteins such as myosin light chains and calmodulin. In contrast, the neck of PfMyoA potentially contains two degenerate IQ motifs. At present, there is only one light chain known to bind to this region of *P. falciparum*, called <u>m</u>yosin <u>t</u>ail <u>i</u>nteracting <u>p</u>rotein (MTIP). The last 15 amino acids of the heavy chain are sufficient to bind MTIP (3), and MTIP has been crystallized in several conformations bound to this peptide (4, 5). The nomenclature for MTIP derived from the idea that it bound to the “tail” of the molecule, a region which typically acts as a targeting and/or a dimerizing domain. One model of glideosome organization proposes that PfMyoA is anchored via the N-terminal extension of MTIP to integral membrane proteins (<u>g</u>lideosome-<u>a</u>ssociated proteins, or GAP) located in a double-membraned flattened complex called the inner membrane complex, which lies ~25nm below the plasma membrane (reviewed in (2)). In that capacity, MTIP functionally acts like a “tail” in addition to binding and stabilizing the α-helical heavy chain following the motor domain.

Here we express functional PfMyoA using the baculovirus/*Sf*9 cell system, and show that proper folding of the motor domain requires that the heavy chain is co-expressed with a UCS family (UNC-45/CRO1/She4p) myosin co-chaperone from *Plasmodium spp*. UCS family myosin co-chaperones have three domains, an N-terminal tetratricopeptide repeat that binds to HSP-90, a central domain of unknown function, and a C-terminal UCS domain that binds myosin (reviewed in (6)). While many myosins can be folded by endogenous *Sf*9 cell chaperones, the divergent PfMyoA requires a *Plasmodium spp*. specific UCS family chaperone. We also show that PfMyoA binds two light chains, MTIP and a novel essential type light chain (PfELC), identified by mass spectrometry of bands associated with FLAG-tagged PfMyoA purified from *Plasmodium* lysates. PfMyoA containing both light chains moves actin at ~3.8 μm/sec in an *in vitro* motility assay, consistent with the light chain binding region acting as a lever arm that amplifies small nucleotide-dependent motions in the motor domain. The ability to express milligram amounts of PfMyoA in a heterologous expression system allows the motor that powers the invasion of the malaria parasite into vertebrate host cells to be characterized, and is the first step toward reconstituting the glideosome *in vitro*.

## Results

### Expression of PfMyoA in Sf9 cells requires co-expression with a *Plasmodium* chaperone

We expressed two *Plasmodium falciparum* class XIV myosin heavy chain constructs using the baculovirus/*Sf*9 insect cell expression system. One construct was the motor domain (PfMD), which contains the actin-binding and active site. It consisted of heavy chain amino acids 1- Lys768, followed by a FLAG tag to facilitate purification by affinity chromatography. The second construct was the full-length PfMyoA that included the light chain binding domain, followed by a biotin tag for attachment to neutravidin coated-surfaces for *in vitro* motility assays, and a C-terminal FLAG tag for affinity purification. The full-length construct was expressed in combination with the known light chain MTIP, which interacts with the C-terminal 15 amino acids of the PfMyoA heavy chain (3).

We previously showed (7) that expression of soluble, functional class XIV myosin from *Toxoplasma gondii* in *Sf9* cells required co-expression with TgUNC, a myosin co-chaperone of the UCS protein family from *T. gondii*. Similarly, when *Sf9* cells were infected with virus encoding the PfMyoA heavy chain and MTIP, the heavy chain was expressed but was not soluble following cell lysis. We next co-expressed the FLAG-tagged full-length PfMyoA heavy chain, untagged MTIP, and TgUNC chaperone. Soluble protein was obtained, but a considerable amount of TgUNC remained bound to the PfMyoA following purification on a FLAG affinity resin (Fig. 1A). Because chaperones bind only to unfolded proteins and are released once the protein is folded, this result indicated that TgUNC was not fulfilling the same role in folding that it did for TgMyoA (7). We speculated that the homologous chaperone from *Plasmodium spp*. was required.

**Fig. 1.**
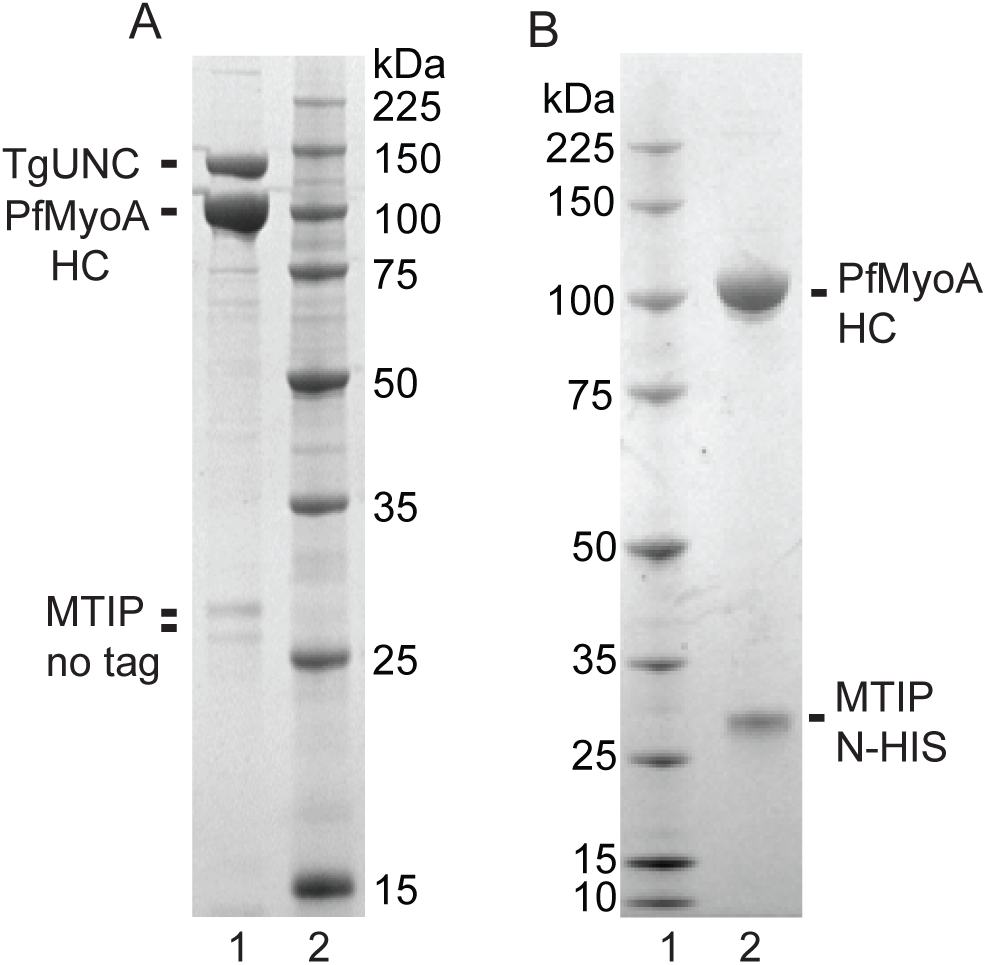
SDS-gels of affinity-purified PfMyoA. (A) The full-length PfMyoA heavy chain was co-expressed with MTIP and TgUNC, the myosin co-chaperone from *T. gondii* (7), and purified on a FLAG-affinity column. Note that the chaperone, TgUNC, co-purifies with the myosin, indicating that some of the protein is misfolded. HC, heavy chain. 12% SDS-gel (B) The full-length PfMyoA heavy chain was co-expressed with MTIP and the putative *Plasmodium* myosin co-chaperone, PUNC. In contrast to panel (A), the chaperone did not co-purify with the myosin, indicating a properly folded protein. 4-12% SDS- gel.

A BLAST search using TgUNC as input identified a putative tetratricopeptide repeat family protein, which was ~90 amino acids longer than most other UNC proteins, which could be the *P. falciparum* myosin co-chaperone (accession number PF3D7_1420200, nomenclature according to http://PlasmoDB.org, or GenBank XP_001348369.1). Our first attempts at expression of the putative chaperone by itself, with native *Plasmodium* codons, resulted in no detectable quantity of protein being expressed in *Sf9* cells. Expressing proteins from *Plasmodium spp*. in heterologous cells is challenging due to its extremely AT-rich genome, which results in large differences from *Sf*9 cell codon preference (8). Another issue is that regions of the putative chaperone contained atypically long stretches of asparagine. To circumvent the latter problem, potential chaperone protein sequences from eight different *Plasmodium spp*. were aligned, and conserved areas identified. A chimeric protein was designed by deleting or changing *P. falciparum* coding sequence to that of *P. knowlesi* in regions where the *P. falciparum* sequence was significantly different from the consensus sequence of the other seven species. To address preferred codon usage, the chimeric chaperone was synthesized using *Sf*9 cell preferred codons. We refer to the putative chimeric chaperone as PUNC (*<u>P</u>lasmodium spp*. UNC).

When PUNC was co-expressed with FLAG-tagged full-length PfMyoA heavy chain and N-HIS tagged MTIP, not only was PfMyoA expressed and soluble, but the resulting myosin was purified free of the PUNC chaperone, consistent with a fully folded protein which releases the chaperone (Fig. 1B). Sedimentation velocity measurements in the analytical ultracentrifuge showed that PfMyoA-MTIP sedimented as a single symmetrical boundary with a sedimentation coefficient of 5.7 ± 0.02S (Fig. 2B). This result verifies the homogeneity of the expressed protein in solution, and is strong evidence for proper folding of the myosin by PUNC. Soluble protein was also obtained when the shorter PfMD construct was co-expressed with PUNC. Affinity-purified MD also showed a single, symmetrical peak, and was characterized by a sedimentation coefficient of 5.0 ± 0.01S (Fig. 2B). The lower S value is consistent with the lower molecular weight and lower asymmetry of MD.

**Fig. 2.**
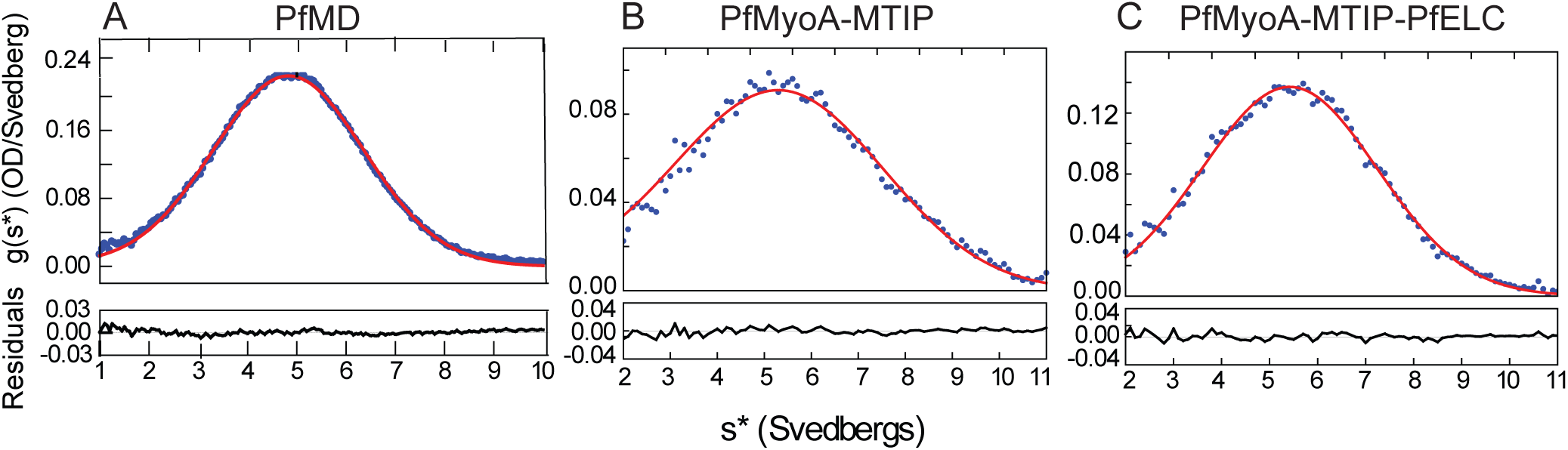
Analytical sedimentation velocity of expressed myosin constructs. All three constructs showed a single, symmetrical peak indicating protein homogeneity. (A) A truncated heavy chain motor domain (MD) construct, PfMD, co-expressed with PUNC sedimented at 5.0 ± 0.01S. (B) Full-length PfMyoA heavy chain co-expressed with PUNC and MTIP sedimented at 5.7 ± 0.02S. (C) Full-length PfMyoA heavy chain co-expressed with PUNC, MTIP, and the newly identified PfELC sedimented at 5.7 ± 0.01S. Solvent conditions: 10 mM HEPES, pH 7.5, 0.15 M NaCl (0.1 M NaCl for PfMD), 1 mM DTT, 20°C. OD, optical density.

An unusual feature of MTIP was that it showed a single band on SDS-gels only when it was tagged at the N-terminus, whether expressed in *Sf*9 cells or in *E.coli*. Expression with untagged or C-terminally tagged MTIP resulted in multiple gel bands, such as seen in Fig. 1A. MTIP immunoprecipitated from *Plasmodium* lysates has also been previously shown to run as multiple bands on SDS-gels, and it was suggested that this may result from various phosphorylation states (9). Mass spectrometry data of N-HIS-MTIP expressed in *Sf*9 cells, however, showed that it was phosphorylated (see later section), yet migrated as a single band. The multiple bands may thus result from proteolytic cleavage at the N-terminus, which is suppressed by the 6xHIS tag.

### *In vitro motility* of PfMyoA

The function of expressed PfMyoA was tested by an *in vitro* motility assay, where surface-bound motors propel rhodamine-phalloidin labeled actin in solution. PfMyoA was specifically attached by its C-terminal biotin tag to a neutravidin-coated coverslip to ensure that all motor domains were available to interact with skeletal actin. When the full-length PfMyoA heavy chain was expressed without light chains, it moved actin at a speed of 0.67 ± 0.16 μm/sec (n=1008). When the heavy chain was co-expressed with MTIP, the speed increased to 1.75 ± 0.32 μm/sec (n=3242) (Fig. 3A). Speeds were analyzed using a semi-automated filament tracking program (see Materials and Methods) and fit to a Gaussian curve. Use of this program permitted evaluation of a large number of filaments without selection bias by the user. The observation that the speed of *in vitro* motility increased in the presence of MTIP is consistent with the idea that speed is proportional to the effective length of the lever arm, which depends on the number of light chains bound to a given myosin (reviewed in (10)).

**Fig. 3.**
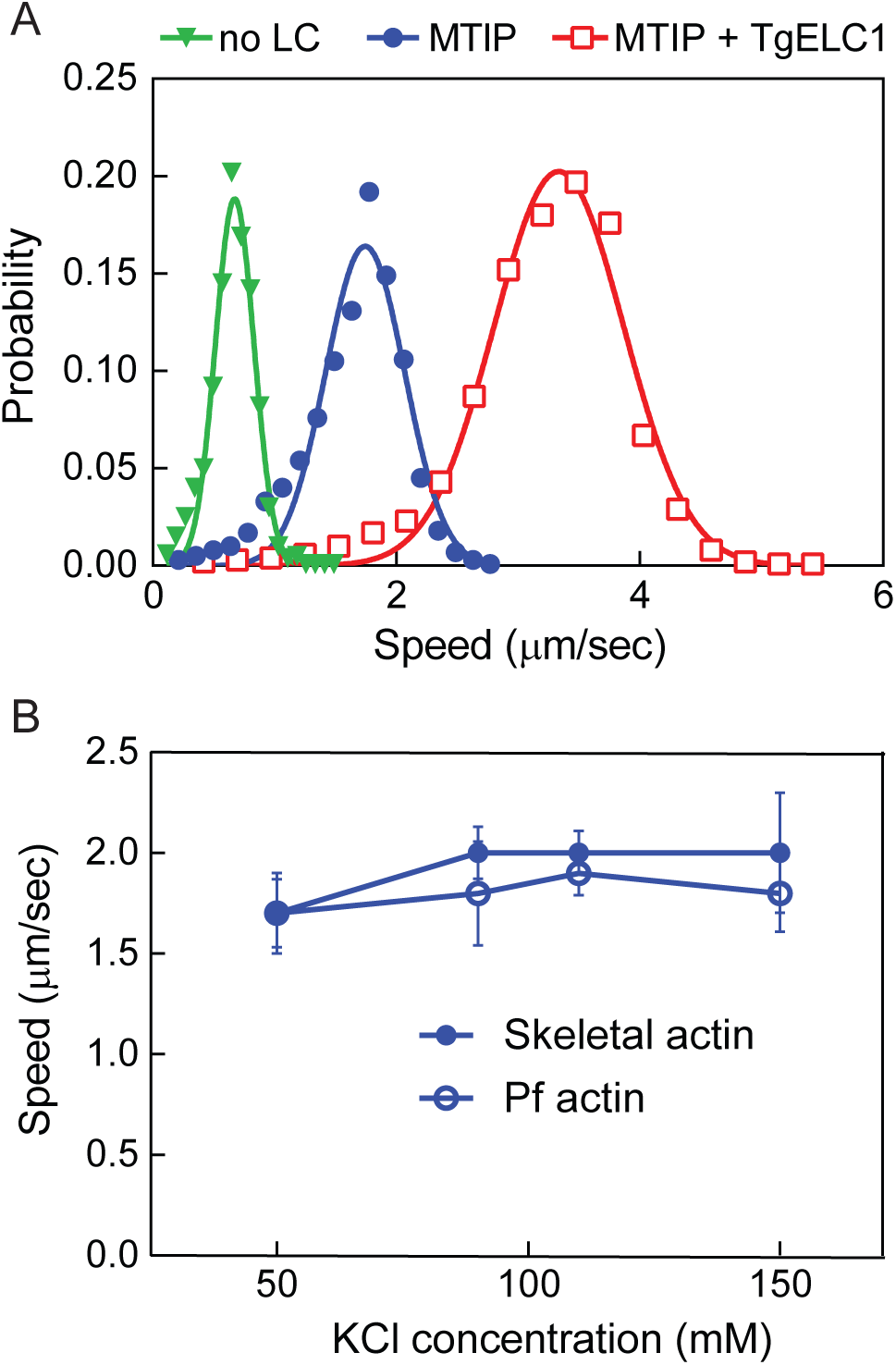
*In vitro* motility. (A) Speeds at which skeletal actin is propelled by full-length PfMyoA containing different bound light chains. Heavy chain only (inverted green triangles), 0.67 ± 0.16 μm/sec (n=1008). Heavy chain and MTIP (filled blue circles), 1.75 ± 0.32 μm/sec (n=3242). Heavy chain, MTIP, and the essential light chain from *T. gondii* (TgELC) (open red squares), 3.3 ± 0.53 μm/sec (n=1691). Conditions: 50 mM KCl, 25 mM imidazole, pH 7.5, 1 mM EGTA, 4 mM MgCl_2_, 10 mM DTT, 2 mM MgATP 30°C. (B) Speed as a function of KCl concentration, using either skeletal actin (filled blue circles) or expressed Plasmodium actin (open blue circles). Conditions: varying KCl, 25 mM imidazole, pH 7.5, 1 mM EGTA, 4 mM MgCl_2_, 10 mM DTT, 2 mM MgATP, 30°C. Data from two independent experiments were pooled.

The speed at which PfMyoA-MTIP moved actin was the same whether we used skeletal muscle actin purified from tissue, or *P. falciparum* actin that was expressed using the baculovirus/*Sf*9 insect cell expression system (see Materials and Methods). Motility speed was similar for both actins over a range of KCl concentration from 50-150 mM KCl (Fig. 3B).

To examine whether the long N-terminal extension present in MTIP affects the speed at which the motor moves actin, we cloned a truncated version of MTIP that lacked the first 60 residues (ΔN-MTIP) for co-expression with the heavy chain. Some light chain extensions interact with actin and slow the speed of the myosin (11). In a direct comparison, the speed at which actin was moved by the construct containing ΔN-MTIP (1.4 ± 0.25 μm/sec, n=1790) was similar to that of the construct containing full-length MTIP (1.3 ± 0.25 μm/sec, n=1978). This result is consistent with the role of the N-terminal extension of MTIP functioning as an adapter for binding GAP45 (12), rather than binding to actin.

### Phosphorylation state of expressed PfMyoA

To identify whether PfMyoA expressed in *Sf*9 cells was phosphorylated, samples were analyzed by a phosphoprotein gel stain. PfMyoA was co-expressed with either the full length MTIP, or ΔN-MTIP. Protein samples were run on an SDS gel and then stained with ProQ Diamond phosphoprotein gel stain (Fig. 4A). The results showed that both the PfMyoA heavy chain and MTIP were phosphorylated. In contrast, the preparation containing the truncated ΔN-MTIP showed little or no MTIP phosphorylation, establishing that the primary site(s) of phosphorylation on this light chain are in the first 60 amino acids.

Once it was determined that both the heavy chain and MTIP were phosphorylated, the subunits were further analyzed by mass spectrometry. The site phosphorylated on the heavy chain was Ser19. On MTIP, Ser55, 58 and 61 were shown to be phosphorylated in two independent protein preparations, with mono-, di- and tri-phosphopeptides identified. In one of the preparations, Ser51 was also phosphorylated. To assess whether these phosphorylation sites impacted the speed of *in vitro* motility, PfMyoA-MTIP was dephosphorylated using calf intestinal alkaline phosphatase (CIP). The efficacy of the dephosphorylation was confirmed using the phosphoprotein gel stain (Fig. 4B), which showed no staining for MTIP and a weak band for the heavy chain. Mass spectrometry confirmed that phosphorylation of Ser19 of the PfMyoA heavy chain decreased from >90% to <9% following dephosphorylation. Dephosphorylation of both the heavy and light chain had no effect on speed in the unloaded *in vitro* motility assay. Speeds of actin movement by PfMyo-MTIP were 1.79 ± 0.3 μm/sec (n=368) as isolated, and 1.73 ± 0.3 μm/sec (n=352) following dephosphorylation. To test if the phosphorylated versus dephosphorylated myosin had large differences in affinity for actin, the *in vitro* motility assay was performed at various salt conditions (15-70 mM) in the absence of methylcellulose, conditions which allow actin to diffuse away from the coverslip surface if not bound to myosin. No differences were seen between the two phosphorylation states.

**Fig. 4.**
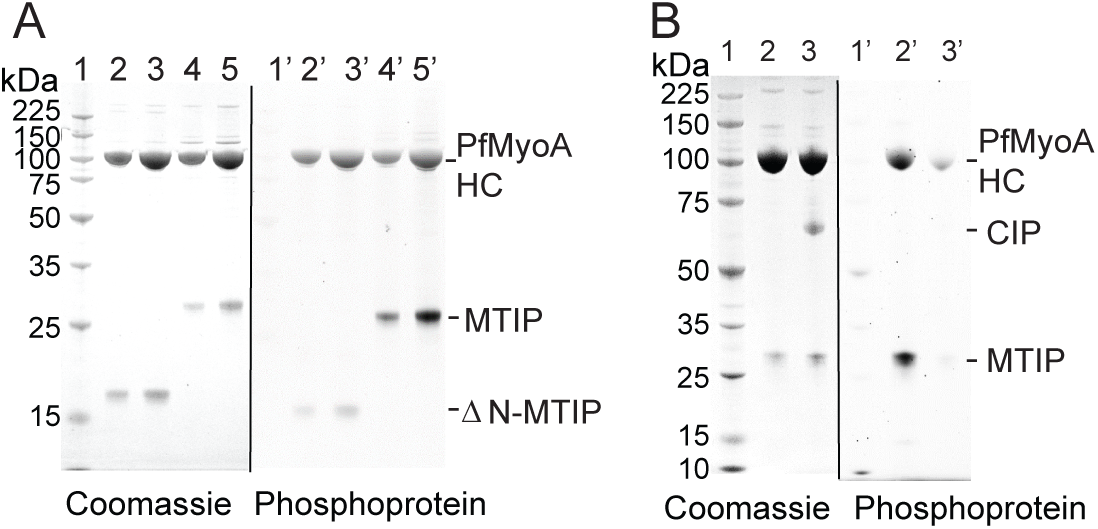
Phosphorylation state of expressed PfMyoA-MTIP. (A) Preparations of PfMyoA-MTIP and PfMyoA-ΔN-MTIP run on 12% SDS-gels and stained with (right panel) ProQ Diamond phosphoprotein gel stain and subsequently with (left panel) Coomassie. Lanes 1,1’, molecular mass markers; lanes 2, 2’ and 3, 3’, two loads of PfMyoA-ΔN-MTIP; lanes 4,4’ and 5,5’, two loads of PfMyoA-MTIP. The heavy chain and full-length MTIP were phosphorylated, but ΔN-MTIP was not. (B) PfMyoA-MTIP run on 4-12% SDS-gels and stained with (right panel) ProQ Diamond phosphoprotein gel stain and subsequently with (left panel) Coomassie. Lanes 1,1’, molecular mass markers; lanes 2,2’ PfMyoA-MTIP as expressed in *Sf*9 cells; lanes 3,3’ following dephosphorylation with calf intestinal phosphatase (CIP). Both the heavy chain and MTIP were dephosphorylated by CIP.

### Effect of phospho-mimic residues at the TEDS site

PfMyoA has a threonine (T417) in the TEDS rule site on the heavy chain (13). This residue fits the pattern seen with most myosins, which either have an acidic residue (D or E) at that site, or a residue capable of being phosphorylated (T or S) to potentially regulate activity by phosphorylation. We engineered a phospho-null T417A mutation, and a phospho-mimic T417D mutation at this site in the PfMD construct. Because this construct without the lever arm is not suitable for *in vitro* motility, we performed actin-activated ATPase assays on MD preparations that contained the mutated heavy chain, and compared it to values obtained with wild-type PfMD. ATPase activity as a function of actin concentration (Fig. 5A), showed that the V_max_ was similar for all three constructs (31.7 ± 1.7 sec^−1^ for WT, 29.1 ± 0.8 sec^−1^ for T417A, 29.7 ± 0.9 sec^−1^ for T417D). A difference was found, however, in the actin concentration at half-maximal activity (K_m_), which reflects the affinity of the motor for actin. Both the WT PfMD and the T417A mutant had similar K_m_ values (13 ± 1.8 μΜ for WT, and 9.8 ± 2.9 μΜ for T417A), consistent with mass spectrometry data showing that Thr417 of PfMyoA was not phosphorylated during expression in *Sf*9 cells. In contrast, the K_m_ of the phospho-mimic T417D was at least two-fold lower (4.4 ± 0.5 μM), indicating a higher affinity for actin. Actin-activated ATPase activity of the full-length PfMyoA-MTIP under the same solvent conditions showed a higher V_max_ (132 ± 6 sec^−1^) and a higher K_m_ (33 ± 4 μM) compared with PfMD (Fig. 5B).

**Fig. 5.**
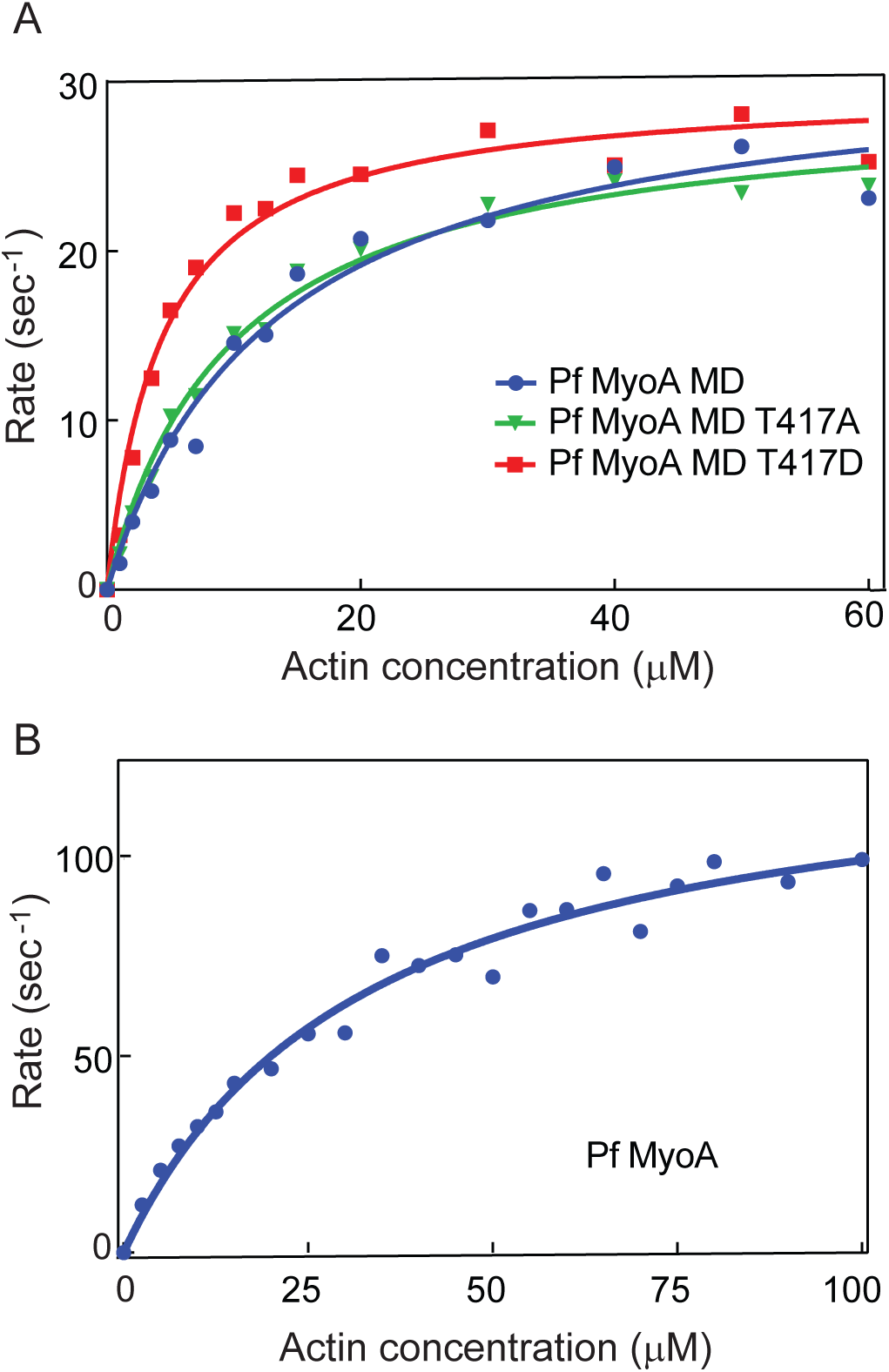
Actin-activated ATPase activity. (A) Plots of ATPase rate versus actin concentration for wild-type PfMyoA MD (filled blue circles), MD with a T417A mutation (green inverted triangles), and MD with a T417D mutation (red filled squares). Data were fit to the Michaelis-Menten equation to yield V_max_ values of 31.7 ± 1.7 sec^−1^ for WT, 29.1 ± 0.8 sec^−1^ for T417A, and 29.7 ± 0.9 sec^−1^ for T417D. K_m_ values were13 ± 1.8 μM for WT, 9.8 ± 2.9 μM for T417A, and, 4.4 ± 0.5 μM for T417D. Conditions: 10 mM imidazole, 5 mM NaCl, 1 mM MgCl_2_, 2 mM MgATP, 1 mM NaN_3_, 1 mM DTT at 30°C. MD WT data were pooled from two independent protein preparations, and four independent experiments. The mutant data were obtained from a single protein preparation. (B) Plots of ATPase rate versus actin concentration for full-length PfMyoA-MTIP. Data were fit to the Michaelis-Menten equation to yield V_max_ =132 ± 6 sec^−1^ and K_m_= 33 ± 4 μM. Conditions: 10 mM imidazole, 5 mM NaCl, 1 mM MgCl_2_, 2 mM MgATP, 1 mM NaN_3_, 1 mM DTT at 30°C. Data from two independent preparations are pooled.

### PfMyoA can bind an essential type light chain in addition to MTIP

We speculated that PfMyoA may bind a second light chain, based on the observation that the class XIV MyoA from *Toxoplasma gondii* binds an essential type light chain (ELC) in addition to MLC1, the homolog of MTIP (7, 14, 15). For proof of principle, PfMyoA heavy chains, MTIP, and TgELC were co-expressed. The purified protein showed three bands at the expected sizes for PfMyoA heavy chain, MTIP and TgELC on SDS gels (Fig. 6A). Furthermore, *in vitro* motility speeds were doubled from ~1.7 μm/sec with MTIP alone, to 3.3 ± 0.53 μm/sec (n=1691) when co-expressed with TgELC (Fig. 3A, red open squares). The same increase in speed was observed when TgELC was added to PfMyoA-MTIP in the final motility buffer, confirming that heavy chain co-expressed only with MTIP remains competent to rebind an extra light chain. This result strongly suggests that PfMyoA binds an essential type light chain in addition to MTIP.

**Fig. 6.**
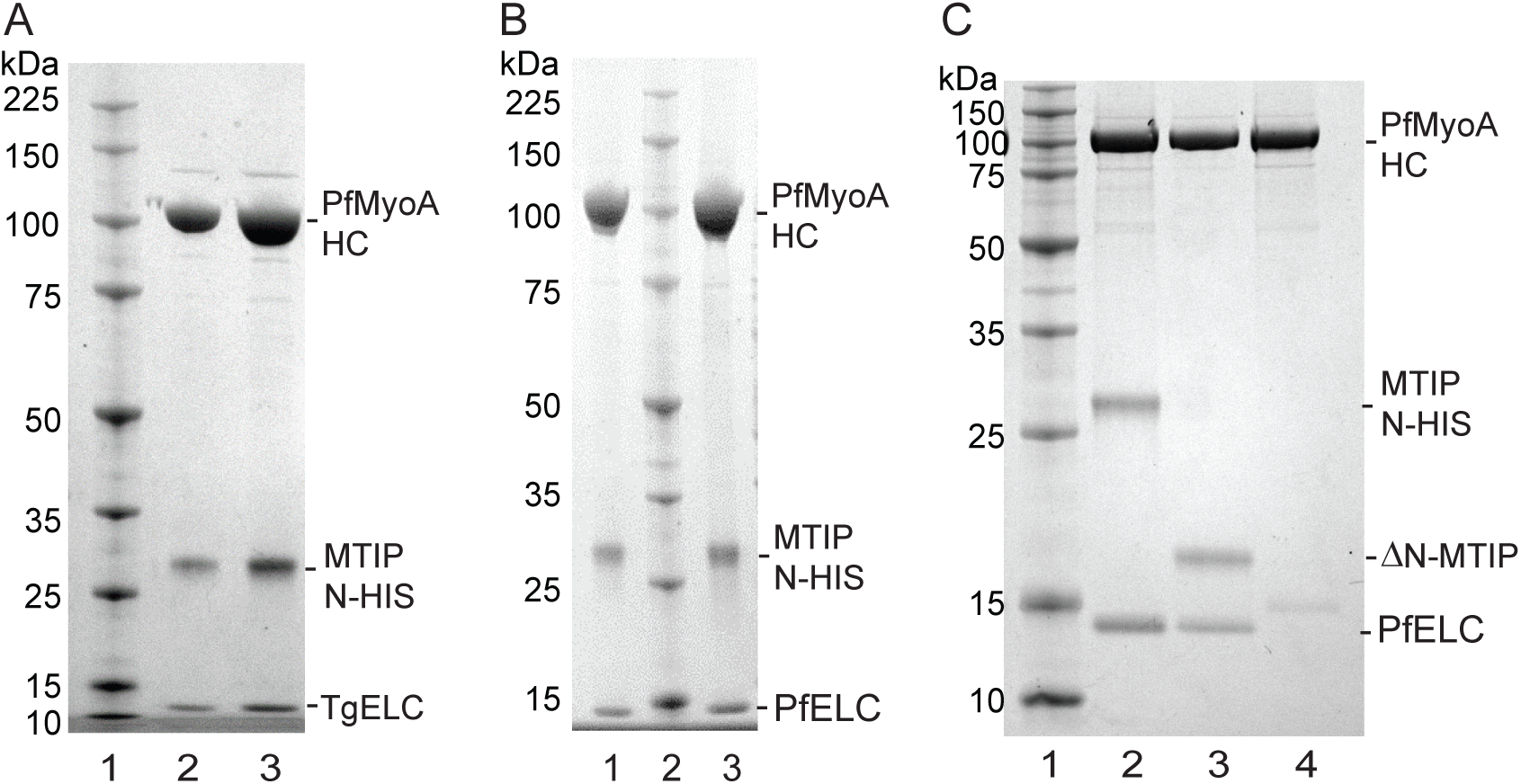
Co-expression of PfMyoA-MTIP with an essential type light chain, and SDS-gel analysis following FLAG-affinity chromatography. (A) Lane 1, molecular mass markers. Lanes 2 and 3, two loads of PfMyoA heavy chain co-expressed with PUNC, MTIP, and TgELC. 4-12% SDS-gel. (B) Lanes 1 and 3, two loads of PfMyoA heavy chain expressed with PUNC, MTIP, and the newly identified PfELC (PF3D7_1017500, GenBank accession number XP_001347455.1). Lane 2, molecular mass markers. 4-12% SDS-gel. (C) Binding of PfELC requires the presence of MTIP. Lane 1, molecular mass markers. PfMyoA heavy chain and PUNC were co-expressed with either: lane 2, PfELC and MTIP; lane 3, PfELC and ΔN-MTIP; or lane 4, PfELC alone, in which case the PfELC did not bind. 12% SDS-gel.

### Search for the *Plasmodium falciparum* essential light chain

To identify potential ELCs from *P. falciparum*, a BLAST search was performed using TgELC as input (Table 1). Four candidate light chains were expressed in bacteria so that the light chain could be added to the final motility buffer of an *in vitro* motility assay. Recombinant virus was also produced for co-expression with the PfMyoA heavy chain in *Sf*9 cells. The highest BLAST score was for *P. falciparum* calmodulin (PfCaM, PF3D7_1434200/ GenBank XP_001348497, 45% identity with TgELC). Calmodulin is the “light chain” for a number of classes of unconventional vertebrate myosins, e.g. class V and class VI. Co-expression of PfMyoA heavy chain and MTIP with PfCaM in *Sf*9 cells showed no PfCaM bound to the purified myosin. Moreover, addition of PfCaM to the final motility buffer showed no enhancement in speed, either in the presence or absence of calcium (Table 1), in contrast to the doubling of speed observed upon addition of TgELC. Other candidate proteins whose functional impact was assayed by their addition to the final motility buffer of an *in vitro* motility assay include (Table 1): (1) PF3D7_1418300 (GenBank accession number XP_001348354.1, 26% identity with TgELC, annotated as a putative calmodulin). It stopped motility when added to the final motility buffer. This putative light chain showed only limited solubility following renaturation after GuHCl denaturation, a property not observed with typical light chains. PF3D7_1418300 also did not co-purify with the PfMyoA heavy chain and MTIP when co-expressed in *Sf*9 cells. (2) PF3D7_0627200 (GenBank accession number XP_966255.2, 21% identity with TgELC, annotated as a putative myosin light chain). This protein slowed motility in the absence of calcium and even further in the presence of calcium. It did not, however, co-purify with the FLAG-tagged PfMyoA heavy chain and MTIP when it was co-expressed with heavy chain in *Sf*9 cells, implying that its binding affinity for the heavy chain was weak. (3) PF3D7_0107000 (GenBank accession number SBT75303.1, 35% identity with TgELC, annotated as a putative centrin-1). It did not enhance speed in the motility assay.

**Table 1.**
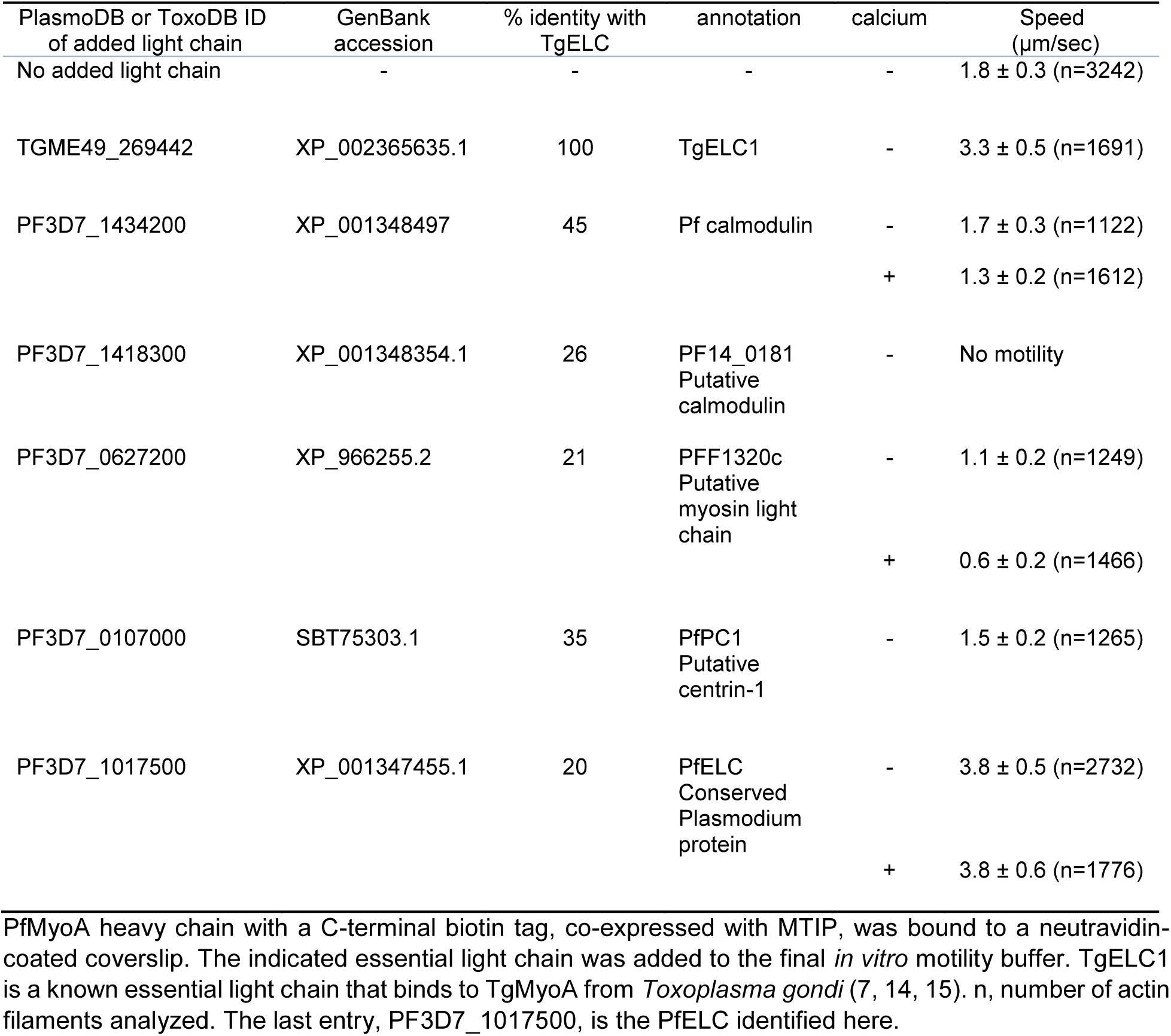
*In vitro* motility speeds of PfMyoA-MTIP to which various putative light chains were added.

All four candidates we tested thus failed to meet our functional criteria of a *bona fide* PfELC, namely that speed should increase when it was added to PfMyoA-MTIP in an *in vitro* motility assay, and it should co-purify with the PfMyoA heavy chain and MTIP when co-expressed in *Sf*9 cells.

### Identification of the *Plasmodium falciparum* essential light chain

To assess whether an additional light chain could be identified directly from the parasite, the PfMyoA heavy chain was tagged in *P. falciparum* using CRISPR/CAS9 technology (16), to introduce a C-terminal dual FLAG tag and a double c-Myc epitope tag (Fig. 7A). After parasite cloning, insertion of the 2c-Myc-2FLAG-tag sequence into the 3’ end of the PfMyoA gene locus was confirmed via PCR analysis (Fig. 7B). Expression of the C-terminal tag in *P. falciparum* schizont stage parasites was verified via Western blot (Fig. 7C) and an immunofluorescence assay using antibodies against the c-Myc epitope (Fig. 7D). Large scale cultures were grown and used to purify PfMyoA (Fig. 7E). Comparing proteins precipitated by FLAG versus control untagged parasite lysates, a 15 kDa light chain-like protein was identified by mass spectrometry, with some homology to calmodulin proteins, PF3D7_1017500 (GenBank accession number XP_001347455.1). Of note, no other obvious light chain or calmodulin-like proteins were identified that were specifically pulled down in FLAG versus control untagged preparations.

**Fig. 7.**
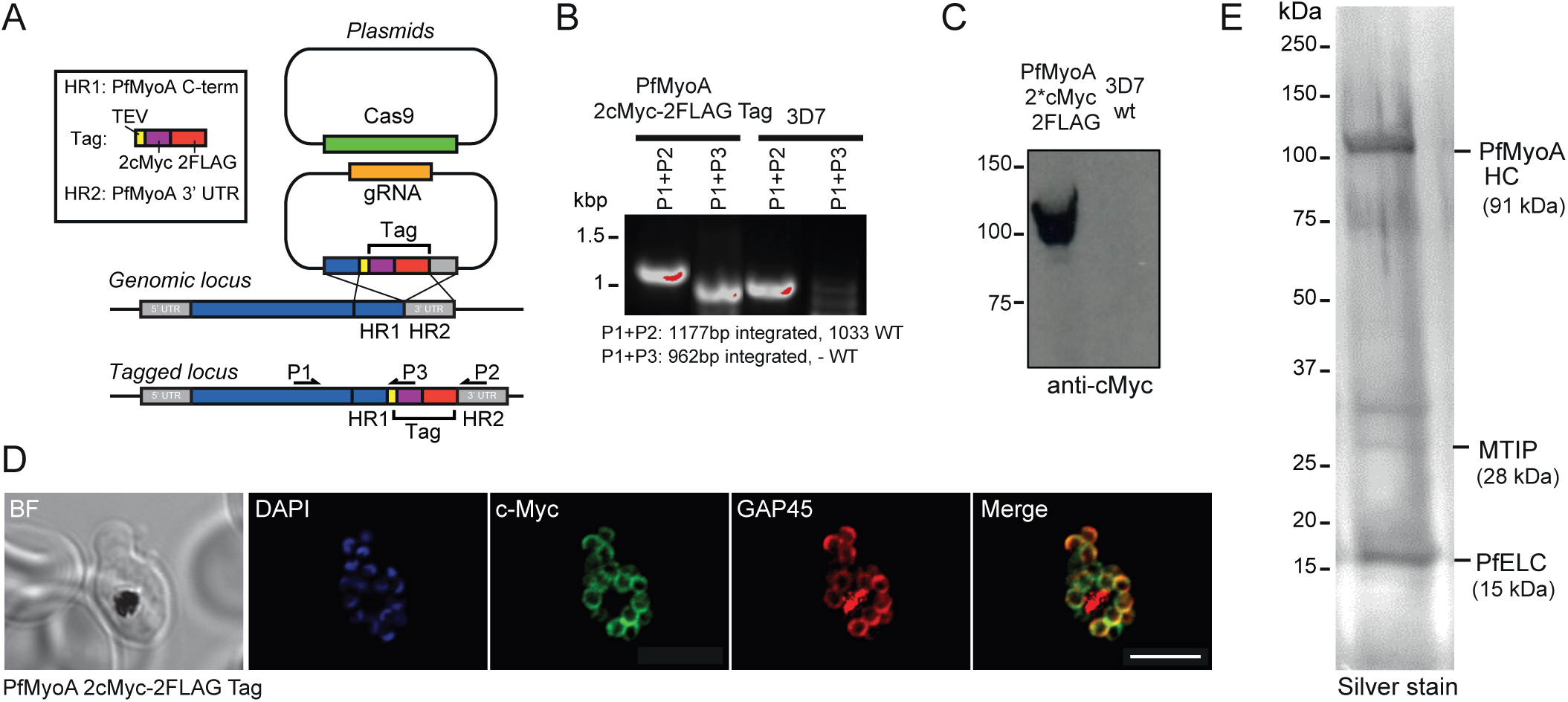
Tagging and localization of the PfELC. (A) A schematic illustrating the strategy used to tag PfMyoA with a C-terminal 2cMyc-FLAG-tag using CRISPR/CAS9. Two plasmids were transfected into *P. falciparum* 3D7 ring stage parasites, one plasmid carrying the guide DNA sequence (AGCTCATATAAGAAAAAAAA) and the desired tag flanked by a fragment of the PfMyoA C-terminus and a fragment of the PfMyoA 3’ UTR (5’ and 3’ homology regions), and a second plasmid containing the cassette for Cas9 expression. Double crossover homologous recombination results in the insertion of the tag sequence between the C-terminal end of PfMyoA and the PfMyoA 3’ UTR. Primers used for detection of the integration of the tag sequence into the genomic loci are shown. (B) PCR analysis of PfMyoA-2cMyc-2FLAG-tag transgenic parasites. Primers able to differentiate between WT sequence and integration of the tag sequence into genomic loci were used to confirm the insertion of the desired tag into the PfMyoA locus. (C) Western blot of PfMyoA-2cMyc-2FLAG-tag transgenic parasites probed with anti-cMyc versus control untagged 3D7 line. (D) Immunofluorescence assay using anti-c-Myc (green), anti-GAP45 (red) and DAPI (blue) in PfMyoA-2cMyc-2FLAG-tag transgenic parasites shows the successful expression of the C-terminal tag and confirmed that the tagged PfMyoA still localised to its expected location in schizonts (at the periphery). BF: bright field. Scale bar: 5 μm. (E) Silver stain on 12 % SDS-PAGE with bands identified as the PfMyoA complex along with the newly identified PfELC.

### Binding and functional effect of PfELC

To assess the ability of PfELC to bind the PfMyoA heavy chain, PfELC was co-expressed in *Sf*9 cells with the PUNC chaperone, PfMyoA heavy chain and MTIP. The purified myosin showed two bound light chains by SDS-gels (Fig. 6B). The purified protein sedimented as a single, homogeneous 5.7 ± 0.01S peak by sedimentation velocity (Fig. 2C). This was the same S value as PfMyoA-MTIP, suggesting that the increase in mass due to PfELC is balanced by an increase in asymmetry due to stabilization of the light chain binding domain by both light chains. Importantly, the *in vitro* motility speed was increased from 1.75 ± 0.32 μm/sec (n=3242) with MTIP only, to 3.8 ± 0.5 μm/sec (n=2732) with PfELC (Fig. 8A and Table 1), similar to the value obtained when TgELC was bound (Fig. 3A and Table 1). There was no difference in the speed of *in vitro* motility when 100 μM calcium was added (3.8 μm/sec ± 0.6, n=1776) (Fig. 8B). Note that 4 mM MgCl_2_ was present in the motility buffer both in the absence or presence of calcium.

**Fig. 8.**
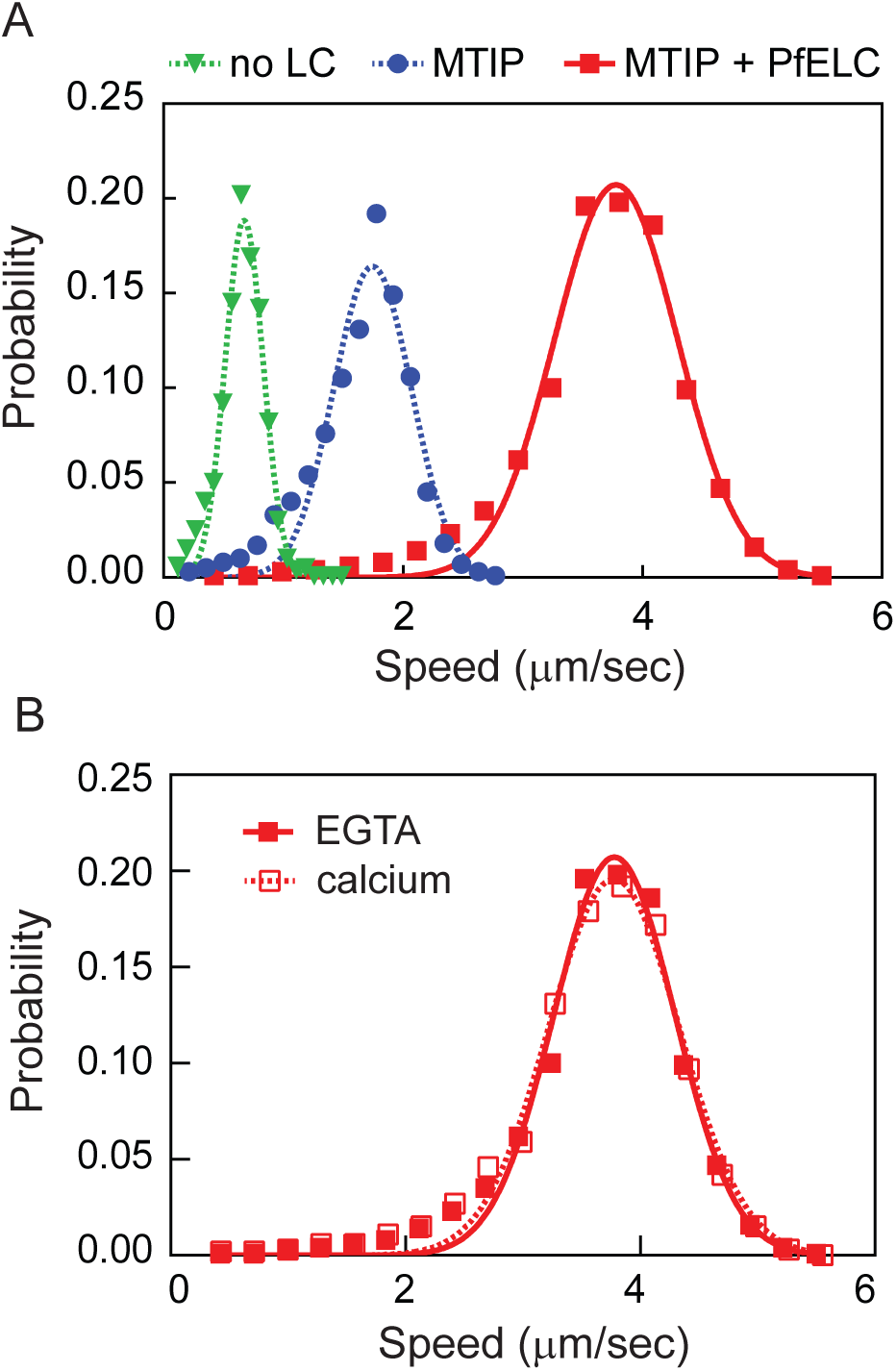
*In vitro* motility of the native PfMyoA with two bound light chains. (A) Dashed lines, speeds of PfMyoA (green inverted triangles) and PfMyoA-MTIP (filled blue circles) repeated from Fig. 3. The filled red squares show the faster speed 3.8 ± 0.5 μm/sec (n=2732) at which actin is propelled when both MTIP and PfELC are bound to the PfMyoA heavy chain. Conditions: 50 mM KCl, 25 mM imidazole, pH 7.5, 1 mM EGTA, 4 mM MgCl_2_, 10 mM DTT, 2 mM MgATP, 30°C. Data from two independent protein preparations were pooled. (B) *In vitro* motility speeds of PfMyoA with both bound light chains were the same in the absence (filled red squares, from panel A) or presence of 100 μM calcium 3.8 ± 0.6 μm/sec (n=1776), (open red squares). Conditions: 50 mM KCl, 25 mM imidazole, pH 7.5, 1 mM EGTA or 100 μM calcium, 4 mM MgCl_2_, 10 mM DTT, 2 mM MgATP, 30°C.

Dephosphorylation of PfMyoA with both light chains bound caused no change in the unloaded *in vitro* motility speed (3.7 ± 0.5 μm/sec, n=765 versus 3.6 ± 0.5 μm/sec, n=808 following dephosphorylation with CIP). PfMyoA with only MTIP bound also showed no change in speed upon dephosphorylation (see earlier section).

The newly identified PfELC, despite its low percent sequence identity with TgELC (20%), appears to be the *bona fide* light chain based on its ability to co-purify with the heavy chain, and double the *in vitro* motility speed compared with the value obtained with MTIP alone. Interestingly, co-expression experiments showed that binding of PfELC to the heavy chain required the presence of MTIP or ΔN-MTIP (Fig. 6C). This result suggests that MTIP enhances the affinity of PfELC for the heavy chain, but the N-terminal extension is not involved. In contrast, MTIP binding does not require PfELC.

## Discussion

Here we show that functional PfMyoA can be expressed in milligram quantities in a heterologous system (*Sf*9 cells) only if a *Plasmodium spp*. myosin co-chaperone of the UCS family is simultaneously co-expressed with the heavy chain. Moreover, the speed at which PfMyoA can move actin in an *in vitro* motility assay is highest (~3.8 μm/s) when two light chains are bound, namely the previously identified heavy chain binding protein MTIP (3, 4), and a novel essential type light chain called PfELC. These results establish that the domain structure of PfMyoA is similar in some respects to more conventional myosins, namely it contains a motor domain that has the actin-binding and active site, and a “neck” that binds light chains, which acts as a lever arm to amplify smaller conformational changes in the motor domain into larger scale movements of actin. Lacking a tail in the more conventional sense, the N-terminal domain of MTIP likely acts as a targeting domain to connect the motor to the inner membrane complex so that relative motion of actin and myosin can occur.

### Chaperone specificity

While most myosins can take advantage of the general cell chaperone present in *Sf*9 cells to properly fold, there have been several instances where this was not the case. We previously showed that the class XIV myosin from *T. gondii*, TgMyoA, required co-expression with a myosin co-chaperone from *T. gondii* called TgUNC (7). TgUNC was not sufficient, however, to fully fold PfMyoA. Chaperones only bind to unfolded proteins, and TgUNC remained bound to PfMyoA following affinity purification. Consequently, we needed to co-express the heavy chain with a UCS family chaperone from *Plasmodium spp*., herein called PUNC. Another example of improper folding of a myosin in *Sf*9 cells by endogenous chaperones was striated muscle myosins (cardiac, skeletal). This failure led to the development of expression of striated muscle myosins in a myocyte cell line C2C12, which by definition has all the factors needed for expression of striated muscle myosin (17). Another example is myosin 15, which is native to inner ear hair cells. To express this myosin in *Sf*9 cells, it needed to be co-expressed with the muscle specific chaperone UNC45b and heat-shock protein 90 (18).

### Phosphorylation sites

The heavy chain and MTIP are phosphorylated during expression in *Sf*9 cells. The site on the heavy chain is Ser19, at the N-terminal region of the motor domain. In *Plasmodium*, protein kinase A (PKA) phosphorylates Ser19 in schizonts prior to egress of merozoites, the stage that infects red blood cells (19). There is also some evidence that cAMP signaling plays a role during sporozoite infection of hepatocytes (20). Interestingly, the other class XIV myosin in *Plasmodium*, PfMyoB, is also phosphorylated at 2 sites near the N-terminus of the heavy chain (Ser16 and Thr17) (19, 21), suggesting a potentially conserved function for phosphorylation at this site.

The other sites phosphorylated during expression in *Sf*9 cells are several residues in the N-terminal extension of MTIP (Ser51, 55, 58 and 61). *In vitro*, both MTIP and glideosome-associated protein 45 (GAP45) can be phosphorylated by calcium-dependent protein kinase 1 (PfCDPK1) (22). The two primary phosphorylation sites by PfCDPK1 were Ser47 and Ser51 of MTIP, with Ser47 the preferred site. We detected no phosphorylation at Ser47 of MTIP when expressed in *Sf*9 cells. *In vivo*, PfCDPK1 is expressed in late schizonts, and was localized to the periphery of merozoites, a location that would allow it to act on the motor complex at the inner membrane complex (22).

When we dephosphorylated both the heavy chain and MTIP, we saw no phosphorylation-dependent difference in the speed at which PfMyoA moved actin in an unloaded *in vitro* motility assay. This was true with only MTIP bound or with both light chains bound. This result does not preclude the possibility that an effect may be seen once this assay is performed under loaded conditions, which more closely mimics the situation that occurs during host cell invasion. Expression of PfMyoA with an N-terminally truncated MTIP lacking 60 amino acids also showed no change in the speed of actin movement in an unloaded *in vitro* motility assay. This result implies that the N-terminal extension of MTIP probably does not bind to actin, which would slow motility (11). It is more likely that the N-terminal extension of MTIP acts primarily as a binding domain to anchor the motor to GAP proteins. The state of phosphorylation of the N-terminus of MTIP may modulate the strength of those interactions.

### TEDS site

PfMyoA lacks an acidic residue in a key actin-binding loop on the heavy chain. This site is referred to as the TEDS rule site, because virtually all myosins have either an acidic residue or a phosphorylatable residue at this position (13). In *Plasmodium* the residue is Thr417, raising the question of whether phosphorylation at this site can regulate interaction of PfMyoA with actin. Proteomic studies to date have not, however, identified this site as phosphorylated in the malaria parasite. Nonetheless, mutation of this residue to the phospho-mimetic Asp in a motor domain construct (T417D) increased the affinity for actin in an ATPase assay, such that at low actin concentrations activity was enhanced compared with the phospho-null. As expected, T417A and the native heavy chain showed superimposable ATPase curves, consistent with this site not being phosphorylated during expression in *Sf*9 cells. In principle, phosphorylation at this site could enhance activity when the effective F-actin concentration is low.

### Interaction of PfMyoA with *Plasmodium* versus skeletal actin

*Plasmodium* actin 1 is markedly divergent from skeletal actin, having only ~80% sequence identity with mammalian actins, a large difference considering the near identity of most actins (23). Although we expected that PfMyoA might interact more optimally with its native actin than with skeletal actin, the speed at which PfMyoA moved both actins in an unloaded motility assay was essentially the same over a range of KCl concentrations (50-150 mM KCl). One caveat is that the *Plasmodium* actin filaments were stabilized by jasplakinolide, and the skeletal actin filaments with phalloidin, which might mask differences that occur with native filaments. At present, however, it has not been possible to maintain stable *Plasmodium* actin filaments in the absence of jasplakinolide. It has been suggested that *Plasmodium* actin is incompletely folded in heterologous systems (24), and that this could explain its aberrant polymerization behavior. Our data do not rule out this possibility, but they do suggest that *Plasmodium* actin expressed in *Sf*9 cells folds properly to the extent that it provides a binding surface for PfMyoA, which can then be moved in a manner indistinguishable from skeletal actin isolated from tissue.

The constant speed seen between 50-150 mM KCl is atypical of vertebrate myosins such as class II smooth muscle myosin or myosin V, which increase in speed with salt concentration (25, 26). This result implies that the interface on PfMyoA that interacts with actin may involve fewer ionic interactions than with these other myosins. Loop 2 of PfMyoA, a positively charged surface loop that is involved in the initial electrostatic interaction with actin, has fewer charged residues and is at least 15 amino acids shorter than either smooth muscle myosin or myosin V.

### Duty ratio estimates

The ability to observe robust motility in the *in vitro* motility assay even at salt concentrations approaching physiological ionic strength is more typically seen with myosins that have a high duty ratio, meaning that the motor spends a large fraction of its ATPase cycle in a state that is strongly bound to actin (26). The duty ratio of PfMyoA can be estimated from the time spent strongly attached to actin relative to the total ATPase cycle time. Assuming a unitary step-size of ~5.5 nm (d_uni_) for PfMyoA (based on 6 nm for single headed class II myosins with two bound light chains (27), or 5.3 nm for TgMyoA (28)), and a measured speed of 3.8 μm/s (v) in the motility assay, yields a strongly attached time (t_on_) of ~1.4 msec (t_on_= d_uni_/v). The maximal actin-activated ATPase activity of ~130 sec^−1^ gives a total cycle time of ~7.7 msec. The estimated duty cycle for PfMyoA is thus ~18%, greater than the 3-4% duty ratios of conventional class II myosins that work in large ensembles because they form filaments, but less than that required for myosins that transport cargo as single motors (minimum 50% duty ratio) (reviewed in (29)). A duty ratio of ~18% implies that even small ensembles of motors (~6) could move actin at maximal speed. The architecture of the glideosome and the labile and potentially short actin filaments may necessitate a higher duty cycle motor.

### Identification of the PfMyoA essential light chain

It is known that the last 15 amino acids of the PfMyoA heavy chain are sufficient to bind MTIP (3). Recent studies with the class XIV myosin from *T. gondii* have identified two essential light chain isoforms that bind mutually exclusively to the heavy chain following the motor domain (14, 15), and which functionally act to promote fast movement of actin (7). Using TgELC1 as input for BLAST searches, there were many false leads for the *bona fide Plasmodium* essential light chain (Table 1). Interestingly, expression of the PfMyoA heavy chain with only MTIP resulted in a stable myosin that was capable of subsequently re-binding bacterially expressed exogenous light chains, including TgELC, suggesting the capacity for a second, native essential light chain. Towards its identification, tagged endogenous PfMyoA was precipitated from large scale parasite cultures. Among bound proteins, a specific binding protein, called here PfELC, was identified, which showed some limited homology to TgELC1 (20% identity) or TgELC2 (21%) (Fig. S1). Secondary and tertiary protein prediction using the RaptorX web portal (http://raptorx.uchicago.edu/) predicted that the protein has seven helices (58% helical), connected by loops (Fig. S2), typical of a light chain that could bind to and stabilize a single α-helical myosin heavy chain.

In support of its association with the glideosome, PfELC was also previously found in a proteomic study of proteins greatly reduced in abundance following selective inhibition of N-myristoylation, which leads to failure to assemble the inner membrane complex incorporating the major glideosome-associated proteins (30). Critically, when expressed as a complex, *in vitro* motility speed doubled comparing PfMyoA and PfMTIP alone versus inclusion of PfELC. Thus, despite its low sequence identity to TgELC, PfELC appears to be a *bona fide* light chain of the PfMyoA-dependent glideosome.

### Effect of calcium

The first isoform of the essential light chain from *T. gondii* (TgELC1) was identified from an analysis of proteins that bound more tightly to TgMyoA in the presence of calcium (14). The residues in TgELC1 predicted to chelate calcium were D15, D17 and D19 (15). In PfELC only one of these three residues (D21 in PfELC) is conserved (Fig. S1); the other two residues are Ser. Alignment of known calcium binding sites in PfCaM with potential motifs in PfELC suggests that PfELC is not capable of binding calcium (Fig. S3). Functionally, we observed no difference in the speed of actin movement with PfMyoA in the presence or absence of 100 μM calcium, with 4 mM MgCl_2_ present in the motility assay buffer. Either PfELC does not bind a divalent cation, or under our buffer conditions the site is occupied by magnesium, which is present at a 40-fold higher concentration than calcium (15). The free calcium concentration in the *Plasmodium* cytoplasm has been measured to be only 350 nM (31). Consistent with our results, an earlier study in which a small amount of PfMyoA was isolated from parasites and analyzed by *in vitro* motility showed no difference in speed between pCa <8 to pCa 4.1 (9). Remarkably, their speed of movement in the motility assay was 3.5 μm/s, very similar to the 3.8 μm/s measured here. In the former study, it was likely that the essential light chain was bound to the heavy chain, but not identified on the SDS-gels used due to its small (~15 kDa) size.

### Conclusions

The speeds we measured under unloaded conditions (~3.8 μm/s) exceed that needed for an ~1μm merozoite to invade a red blood cell in less than 20 sec (0.05 μm/s) (32). The force-velocity curve of myosin shows that speed decreases under load, and thus our measured speeds are compatible with the biology of cellular invasion. Our results solidify the idea that the domain structure of class XIV myosins consists of a motor domain followed by two light chains that bind to the C-terminal region of the heavy chain. The light chain binding domain acts as a lever arm to amplify smaller nucleotide-dependent motions in the motor domain. In the absence of a true “tail” the N-terminal region of MTIP likely plays that role by anchoring the motor into the inner membrane complex via integral GAP proteins. The ability to express PfMyoA in milligram quantities allows needed structural and functional studies to be performed with the motor that powers invasion of host cells to cause malaria, a disease that is a major global health issue. PfMyoA is also a novel druggable target for inhibition of cellular invasion.

## Materials and Methods

Details of *Plasmodium* actin 1 expression, purification, and labeling for visualization in the motility assay, mass spectrometry of PfMyoA expressed in *Sf*9 cells, and mass spectrometry of large scale FLAG-tagged immunoprecipitation of PfMyoA-2cMyc-2FLAG from culture are provided in S1 Materials and Methods.

### Expression Constructs

The full-length PfMyoA heavy chain (PlasmoDB ID PF3D7_1342600/ GenBank accession number XM_001350111.1) was obtained by PCR using gBlocks^®^ Gene Fragments (Integrated DNA Technologies) with *Sf*9 cell preferred codons as template. A 13 amino acid linker (NVSPATVQPAFGS) was added to the C-terminus of the PfMyoA heavy chain to separate it from an 88 amino acid segment of the *Escherichia coli* biotin carboxyl carrier protein (33, 34) that gets biotinated during expression in *Sf*9 cells, followed by a C-terminal FLAG tag. The PCR product was cloned into the baculovirus transfer vector pAcSG2 (BD Biosciences) to make recombinant virus.

A shorter PfMyoA motor domain construct that lacked the light chain binding region was truncated at residue K768, followed by a Gly-Ser linker and a C-terminal FLAG tag. This heavy chain fragment was cloned into baculovirus transfer vector pAcSG2 (BD Biosciences). Two other constructs, in which the TEDS site motif Thr (T417) was mutated to either Ala (phospho-null) or Asp (phospho-mimic) were also cloned.

The PfMTIP gene (PF3D7_1246400/GenBank accession number XM_001350813.1) was PCR-amplified from gBlocks^®^ Gene Fragments using *Sf*9 preferred codons, and cloned with an N-terminal 6xHIS-tag in both pAcSG2 for *Sf*9 cell expression, and pET19 (Novagen) for bacterial expression. In the ΔN-MTIP version, the 60 amino acid N-terminal extension of both MTIP-pAcSG2 and MTIP-pET19 was removed by site directed mutagenesis. The N-HIS tag was retained, and MTIP begins with S61.

Four potential essential light chain candidates were cloned using *Sf*9 preferred codons into a bacterial expression vector, and additionally three of these were cloned in a baculovirus vector. PF3D7_1434200 (GenBank accession number XM_001348461.1) had an N-terminal 6xHIS tag in both pFastBac and pET19. PF3D7_1418300 (GenBank accession number XM_001348318.1) was C-terminally 6xHIS tagged in both pAcSG2 and pET3. PF3D7_0627200 (GenBank accession number XM_961162.2) had an N- terminal HIS tag in pFastBac and no tag in pET3. PF3D7_0107000 (GenBank accession number SBT75303) was only cloned for bacterial expression using pET19 with an N-terminal 6xHIS tag. TgELC used as a control was described in (7).

The PfELC gene (PF3D7_1017500/GenBank accession number XM_001347419.1) identified here was cloned using *Sf*9 preferred codons using gBlocks^®^ Gene Fragments. The resulting PCR product was cloned into the bacterial expression vector pET19 with an N-terminal 6XHIS tag. For expression in *Sf*9 cells, it was cloned without a tag into pFastBac (ThermoFisher Scientific) for recombinant virus production.

The Plasmodium UCS family chaperone, herein called PUNC, is a chimeric clone made using the coding sequence of *P. falciparum* (PF3D7_1420200/GenBank accession number XM_001348333.1) but substituting sequences from *P. knowlesi* (PKNH_1337800/GenBank accession number XM_002260772) in regions that *P. falciparum* did not show consensus among other *Plasmodium* species. Sequence is available upon request. A PCR product was made using gBlocks^®^ Gene Fragments with *Sf*9 cell preferred codons. The resulting chaperone, with a Myc tag at the C- terminus, was cloned into the baculovirus transfer vector pAcSG2 (BD Biosciences) for recombinant virus production. TgUNC used as a control was described in (7).

Plasmodium actin 1 (PF3D7_1246200/GenBank accession number XM_001350811.1) followed by a linker and human thymosin-β4 was synthesized using gBlocks^®^ Gene Fragments with *Sf*9 cell preferred codons, and cloned into pAcUW51 for expression in *Sf*9 cells as described previously (35).

### Myosin Expression and Purification

The full-length PfMyoA heavy chain or the MD was co-expressed with the co-chaperone PUNC in *Sf*9 cells. As indicated, either a 6x-HIS tagged MTIP alone, or both MTIP and an essential type light chain were co-expressed. The cells were grown for 72 h in medium supplemented with 0.2 mg/ml biotin, harvested, and lysed by sonication in 10 mM imidazole, pH 7.4, 0.2 M NaCl, 1 mM EGTA, 5 mM MgCl_2_, 7% (w/v) sucrose, 2 mM DTT, 0.5 mM 4-(2-aminoethyl) benzenesulfonyl fluoride, 5 μg/ml leupeptin, and 2.5 mM MgATP. An additional 2.5 mM MgATP was added prior to clarifying at 200,000 X *g* for 40 min. The supernatant was purified by FLAG-affinity chromatography (Sigma- Aldrich). The column was washed with 10 mM imidazole, pH 7.4, 0.2 M NaCl, and 1 mM EGTA. The myosin was eluted from the column in the same buffer containing 0.1 mg/ml FLAG peptide. The fractions of interest were combined, concentrated with an Amicon centrifugal filter device (Millipore), and dialyzed against 10 mM imidazole, pH 7.4, 0.2 M NaCl, 1 mM EGTA, 55% (v/v) glycerol,1 mM DTT, and 1 μg/ml leupeptin for storage at -20°C.

### Light Chain Expression and Purification

Bacterially expressed MTIP and PfELC HIS-tagged light chains were expressed in BLR(DE3) competent cells and grown in LB broth. Cultures were grown overnight at 27°C following induction with 0.7 mM isopropyl b-D-thiogalactopyranoside, after which the pellet was harvested and frozen. Pellets were lysed by sonication in 10 mM sodium phosphate, pH 7.4, 0.3 M NaCl, 0.5% (v/v) glycerol, 7% (w/v) sucrose, 7 mM β-mercaptoethanol, 0.5 mM 4-(2 aminoethyl)benzenesulfonyl fluoride, and 5 μg/ml leupeptin. The cell lysate was clarified at 26,000 X *g* for 30 min after which the supernatant was boiled for 10 min in a double boiler, clarified at 26,000 X *g* for 30 min, and loaded on a HIS-Select nickel affinity column (Sigma-Aldrich). The resin was washed in buffer A (10 mM sodium phosphate, pH 7.4 and 0.3 M NaCl before eluting with buffer A containing 200 mM imidazole. The protein was concentrated and dialyzed overnight against 10 mM imidazole, pH 7.4, 150 mM NaCl, 1mM EGTA, 1mM MgCl_2_, 1 mM DTT, and 55% (v/v) glycerol and stored at -20°C.

### Sedimentation Velocity

Sedimentation velocity runs were performed at 20°C in an Optima XLI analytical ultracentrifuge (Beckman Coulter) using an An60Ti rotor at 40,000 rpm. The solvent was 10 mM HEPES, pH 7.5, 0.1 or 0.15 M NaCl, and 1 mM DTT. The sedimentation coefficient was determined by curve fitting to one species using the d*c*/d*t* program (36).

### *In Vitro* Motility

A nitrocellulose-coated flow cell was prepared by adding 0.5 mg/ml biotinylated bovine serum albumin, BSA, in buffer A (150 mM KCl, 25 mM imidazole, pH7.5, 1 mM EGTA, 4 mM MgCl_2_, and 10 mM DTT) for 1 min followed by three rinses of 0.5 mg/ml BSA in buffer A. The flow cell was then infused with 50 μg/ml neutravidin (Thermo Scientific) in buffer A for 1 min followed by three rinses with buffer A. Before use, PfMyoA was mixed with a 2-fold molar excess of actin and 5 mM MgATP and spun for 20 min at 350,000×g to remove myosin that was unable to dissociate from actin in the presence of MgATP. PfMyoA (0.5 μM), which had a biotin tag at the C-terminus of the heavy chain, was infused into the neutravidin-coated flow cell and incubated for 1 min, then rinsed 3 times with buffer B (50 mM KCl, 25 mM imidazole, pH 7.5, 1 mM EGTA, 4 mM MgCl_2_, and 10 mM DTT). The chamber was then washed two times with 1 μΜ unlabeled F-actin in buffer B, which was first sheared by vortexing, to block any remaining MgATP-insensitive motors. The flow cell was then washed three times with buffer B containing 2 mM MgATP, followed by three washes with buffer B. Rhodamine-phalloidin-labeled actin was then introduced for 30 sec, followed by one rinse with buffer B, and one rinse with buffer C. Buffer C is buffer B plus 0.5% (wt/vol) methylcellulose, 25 μg/ml PfMTIP, and 25 μg/ml PfELC (or whatever essential light chain was being tested), and oxygen scavengers (50 μg/ml catalase (Sigma-Aldrich), 125 μg/ml glucose oxidase (Sigma-Aldrich), and 3 mg/ml glucose). Lastly, buffer C containing 2 mM MgATP was flowed twice through the flow cell. Assays were performed at 30°C. When assays were performed in the presence of calcium, the light chains were dialyzed against buffer containing 0.1 mM calcium and no EGTA, and 1 mM EGTA in the motility buffers was replaced with 0.1 mM calcium.

Actin movement was observed at 30°C using an inverted microscope (Zeiss Axiovert 10) equipped with epifluorescence, a Rolera MGi Plus digital camera, and dedicated computer with the Nikon NIS Elements software package. Data were analyzed using a semi-automated filament tracking program described previously (37). The speeds of >1000 filaments were determined. Speeds were fit to a Gaussian curve.

### SDS gels

Proteins were separated on either a 12%, or a 4–12% gradient Bis-TrisNuPAGE gel (Invitrogen) and run in MOPs buffer according to the NuPAGE technical guide.

### Dephosphorylation of PfMyoA

30 Units of calf intestinal alkaline phosphatase (10U/μl) NEB was added to 20 μl of 15.4 μM PfMyoA in 100 mM KCl, 25 mM imidazole, pH7.5, 1 mM EGTA, 4 mM MgCl_2_, and 1 mM DTT, and incubated for 30 min at 30°C.

### Phosphoprotein Gel Stain

Protein samples were separated on either a 12%, or a 4–12% Bis-TrisNuPAGE SDS-gel (Invitrogen) run in MOPs buffer according to the NuPAGE technical guide. The gels were first stained with ProQ Diamond phosphoprotein gel stain (Invitrogen) according to the manufacturer’s instructions. The gels were imaged using a Pharos FX Plus scanner (532 excitation, 605 filter) to detect phosphorylated bands. Gels were subsequently stained with Coomassie blue to identify all protein bands.

### Actin-activated ATPase Activity

The actin-activated ATPase activity of PfMyoA was determined at 30°C in 10 mM imidazole, 5 mM NaCl, 1 mM MgCl_2_, 1 mM NaN_3_, 1 mM DTT as a function of skeletal actin concentration. A low salt concentration was needed to keep the K_m_ values as low as possible so that V_max_ could be achieved. PfMyoA-MTIP concentration was 35 nM, and PfMD concentration was 63 nM. The assay was initiated by the addition of 2 mM MgATP and stopped with SDS at 4 time points every min for 4 minutes for PfMyoA-MTIP or every 1.5 minutes for MD. Inorganic phosphate was determined colorimetrically as described in (38). Data were fit to the Michaelis-Menten equation.

### Native Tagging of PfMyoA

A gene fragment coding for a TEV protease cleavage site followed by the c-Myc epitope tag (2x) and a dual FLAG tag, flanked by a fragment of the C-terminal end of PfMyoA and a fragment of the PfMyoA 3’ UTR, was synthesised by GeneART (Thermo Fisher Scientific) and inserted into the pL6-eGFP CRISPR plasmid (16) (Fig. 7). The pL6-eGFP plasmid was linearized for cloning using the SacII/AflII restriction sites. The guideDNA sequence (AGCTCATATAAGAAAAAAAA) was inserted into the same plasmid using the BtgZI adaptor site (16), producing the completed pL7-PfMyoA-2cMyc-2FLAG-tag plasmid. All cloning steps was performed using Gibson Assembly (39). *P. falciparum* 3D7 strain parasites were maintained in RPMI-based media supplemented with O^+^ human erythrocytes at 4% haematocrit and 0.5% AlbuMAX II (Life Technologies), according to established methods (40) with genetic transformation performed as previously described (41). Briefly, ring stage parasites at 8-10% parasitemia were transfected with 60 μg of the pL7-PfMyoA-2cMyc-2FLAG plasmid, along with 60 μg of the pUF1-Cas9 plasmid, which expresses the Cas9 endonuclease (16). Positive drug selection was performed one day post-transfection using 2.5 nM WR99210 and 1.5 μM DSM1 and maintained until stable parasite growth was achieved. The transgenic parasites were then cloned by limiting dilution at concentrations of 0.25 parasites/well, 0.5 parasites/well and 1 parasite/well. Insertion of the tag sequence was determined by PCR analysis of parasite genomic DNA, using primers specific for integration of the tag sequence at the PfMyoA gene locus (Fig.7). Clones with the integrated tag sequence were then maintained in media without drug for further downstream analysis.

### Immunofluorescence Assay of *P. falciparum* Schizonts

Thin smears of *P. falciparum* schizont stage cultures were prepared on glass slides and allowed to air-dry. The smears were fixed in 4% paraformaldehyde/0.01% glutaraldehyde/Phosphate Buffered Saline (PBS) for 1 hour at room temperature. Fixed smears were then permeabilized with 0.1% Triton X-100 for 10 minutes at room temperature and blocked with 3% BSA/PBS for 1 hour at room temperature. Staining was performed with primary rat anti-c-Myc (JAC6, Abcam) diluted 1:200 and primary rabbit anti-GAP45 (42) diluted 1:500, followed by secondary Alexa-Fluor-conjugated antibodies (Life Technologies, 1:500 dilution). All antibody dilutions were performed in 1% BSA/PBS. Confocal images were acquired using a Zeiss LSM 510 Laser scanning confocal microscope.

## Acknowledgements

We thank Kate Heesom and Marieangela Wilson from Bristol University Proteomics Facility for experimental support, and Karen Brack at the University of Vermont for mass spectrometry sample preparation and analysis. This work was supported by National Institutes of Health grants AI117476 to KMT and HL124041to MJP. JB is supported by funding from the Wellcome Trust (Investigator Award 100993/Z/13/Z) and Human Frontier Science Organization (Young Investigator Program Award RGY0066/2016).

## Author contributions

CSB cloned, expressed, and characterized PfMyoA *in vitro;* CLT and RM identified PfELC in parasites; MJP performed mass spectrometry; EBK performed analytical ultracentrifugation and protein expression; PMF expressed *Plasmodium* actin; CSB, JB, and KMT designed research and wrote the paper.

